# Codon usage bias creates a ramp of hydrogen bonding at the 5′-end in prokaryotic ORFeomes

**DOI:** 10.1101/811612

**Authors:** Juan C. Villada, Maria F. Duran, Patrick K. H. Lee

## Abstract

Codon usage bias exerts control over a wide variety of molecular processes. The positioning of synonymous codons within coding sequences (CDSs) dictates protein expression by mechanisms such as local translation efficiency, mRNA Gibbs free energy, and protein co-translational folding. In this work, we explore how codon variants affect the position-dependent content of hydrogen bonding, which in turn influences energy requirements for unwinding double-stranded DNA. By analyzing over 14,000 bacterial, archaeal, and fungal ORFeomes, we found that *Bacteria* and *Archaea* exhibit an exponential ramp of hydrogen bonding at the 5′-end of CDSs, while a similar ramp was not found in *Fungi*. The ramp develops within the first 20 codon positions in prokaryotes, eventually reaching a steady carrying capacity of hydrogen bonding that does not differ from *Fungi*. Selection against uniformity tests proved that selection acts against synonymous codons with high content of hydrogen bonding at the 5′-end of prokaryotic ORFeomes. Overall, this study provides novel insights into the molecular feature of hydrogen bonding that is governed by the genetic code at the 5*′*-end of CDSs. A web-based application to analyze the position-dependent hydrogen bonding of ORFeomes has been developed and is publicly available (https://juanvillada.shinyapps.io/hbonds/).

---

Codon usage controls protein synthesis through a variety of mechanisms^1,2^. A number of classic works have established the links between codon usage and mRNA translation^3–5^, with important insights into the physiological consequences of synonymous mutations^6,7^. The specific arrangement of synonymous codons in coding sequences (CDSs) has been shown to serve as a regulatory mechanism of translation dynamics^8^ and protein co-translational folding^9^. In particular, the 5′-end region of CDSs has strong effects on translation where synonymous codon choice is associated with targeting efficiency of signal peptides^10^, ramp of translation efficiency^11^, local folding energy^12^, modulated protein expression^13^, and recognition of nascent peptides by the signal recognition particle^14^.

Similar to translation, codon usage bias has been associated with transcriptional selection^15^ and optimization of transcription efficiency^16^. Recent reports support the idea that codon variants also define the energy and cellular resources required for transcript biosynthesis^17–20^. However, in contrast to translation, the potential links between position-dependent codon usage bias at the 5′-end of CDSs and transcription optimization have yet to be investigated. During transcription, helicases melt the hydrogen bonds in double-stranded DNA (dsDNA) to expose the single stranded DNA (ssDNA) template sequence, while RNA polymerase produces the RNA molecule^21^. Although the role of helicase can be active or passive^22^, the dsDNA unwinding process requires energy^23^ and successful unwinding of the dsDNA is determinant in preventing abortive transcription and translation initiation^24^. In this work, we explore whether a mechanism to optimize transcription efficiency through codon variants exists so that the energy required to unwind the 5′-end of CDSs is reduced. Our central hypothesis stems from the fact that increased GC content of a gene increases the number of hydrogen bonds in its dsDNA, thereby demanding higher unwinding energy^25^. We hypothesized that the energy requirements for unwinding dsDNA of a CDS could be modulated by controlling the usage of synonymous codons to vary the number of hydrogen bonds.

Here, by analyzing over 14,000 ORFeomes (the set of all CDSs in a genome), we provide genomic evidence that codon usage bias creates an exponential ramp of hydrogen bonding at the 5′-end of CDSs in prokaryotes but not eukaryotes (i.e., *Fungi*). The observed prokaryotic ramp can be a possible molecular mechanism that supports the efficient coupling of transcription and translation at the 5′-end of CDSs. Our evidence suggests that synonymous codon variants can fine-tune the energy required for unwinding dsDNA, providing novel insights into the evolution of molecular traits and the trade-offs between the genetic code and physiology of organisms.

## Results

### An optimization space for hydrogen bonding through codon variants

We began our analysis by categorizing codons according to their hydrogen bond content (Supplemental Figure S1). The number of hydrogen bonds in a codon is directly coupled to the GC content of a codon due to the Watson-Crick base pairing of nucleotides^26^. Each codon can contain six to nine hydrogen bonds but most codons tend to have seven or eight (Supplemental Figure S1A). All degenerate amino acids have choices for codons with different number of hydrogen bonds (Supplemental Figure S1B) and the relative content of hydrogen bonding of a codon can be decreased by 25% according to the synonymous codon choice (Supplemental Figure S1C). The optimization space for hydrogen bonding becomes larger when the position-dependent codon usage bias is considered where the overall and local hydrogen bond composition of a CDS can be fine-tuned by introducing synonymous mutations (Figure 1A).

**Figure 1.**
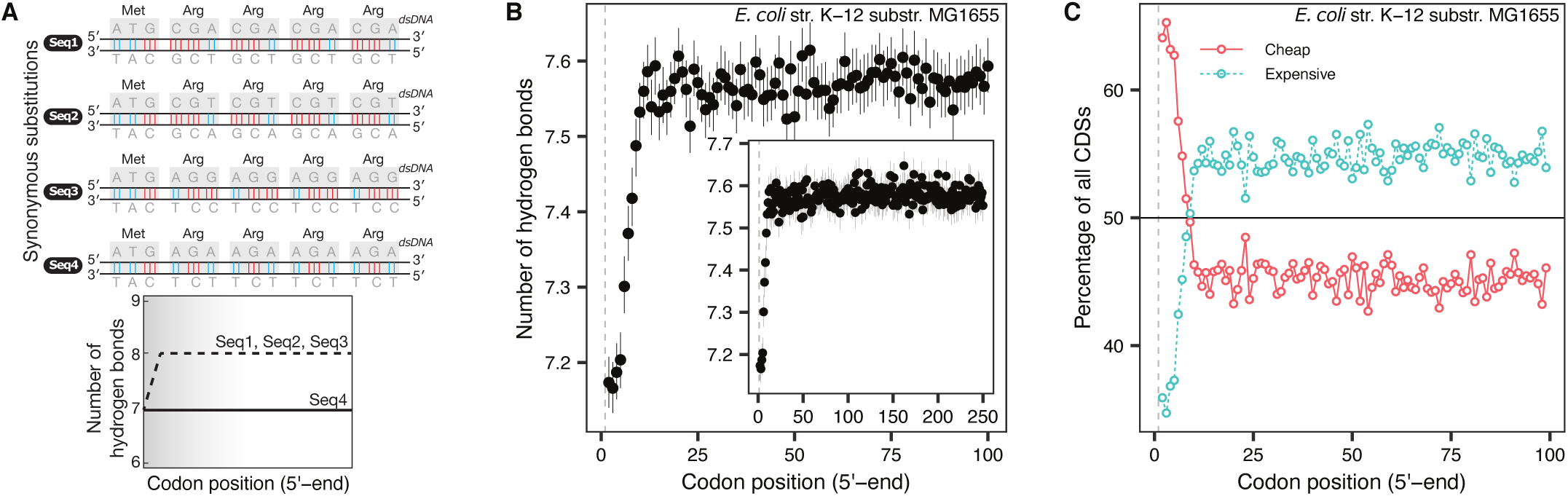
Trade-off between codon usage bias and the number of hydrogen bonds. **(A)** A toy example illustrating synonymous mutations in CDSs can create different distributions of position-dependent hydrogen bonding. **(B)** The number of hydrogen bonds gradually increases in the ORFeome of *E. coli*. The data shown correspond to the mean and 95% confidence interval of the mean with 1,000 bootstraps. The dashed line indicates the position of the start codon. The inset shows the number of hydrogen bonds up to the 250^*th*^ codon position. **(C)** Usage of cheap and expensive codons based on the number of hydrogen bonds along CDSs of *E. coli*.

### Position-dependent content of hydrogen bonding found at the 5′-end of the *E. coli* ORFerome

All CDSs in the ORFeome of *E. coli* were first analyzed to test the hypothesis that the number of hydrogen bonds is position-dependent at the 5′-end. The mean number of hydrogen bonds in each codon position was calculated. We observed that the number of hydrogen bonds per codon gradually increased in a position-dependent manner until about the 15^*th*^ codon position. After this codon position, the number of hydrogen bonds converges to a carrying capacity that remains similar until the 250^*th*^ codon position (Figure 1B). Subsequently, we discretized codons into two groups according to their hydrogen bond content: cheap codons (with six or seven number of hydrogen bonds) and expensive codons (with eight or nine number of bonds). We observed that the group of cheap codons is utilized with high frequency (∼65%) then decreases gradually in a position-dependent manner until reaching an equilibrium at about the 15th codon position (Figure 1C). From the codon position 15th to 100th, the frequency of cheap and expensive codons utilization does not vary by more than ∼5% with cheap codons appearing much less frequently than expensive codons (Figure 1C).

### The ramp of the number of hydrogen bonds at the 5′-end of CDSs is conserved in prokaryotes

Based on the position-dependent arrangement of the number of hydrogen bonds per codon observed in the *E. coli* ORFeome (Figure 1), we fitted three mathematical functions to model the mean number of hydrogen bonds per codon as a function of codon position. According to AIC and BIC criteria, the bounded exponential model with three parameters (initial content, rate, and carrying capacity) produced the best fit (Figure 2A). The fitness of the model showed that the number of hydrogen bonds per codon follows an exponential function of codon position with a positive rate that has a ramp-like shape at the 5′-end of CDSs.

**Figure 2.**
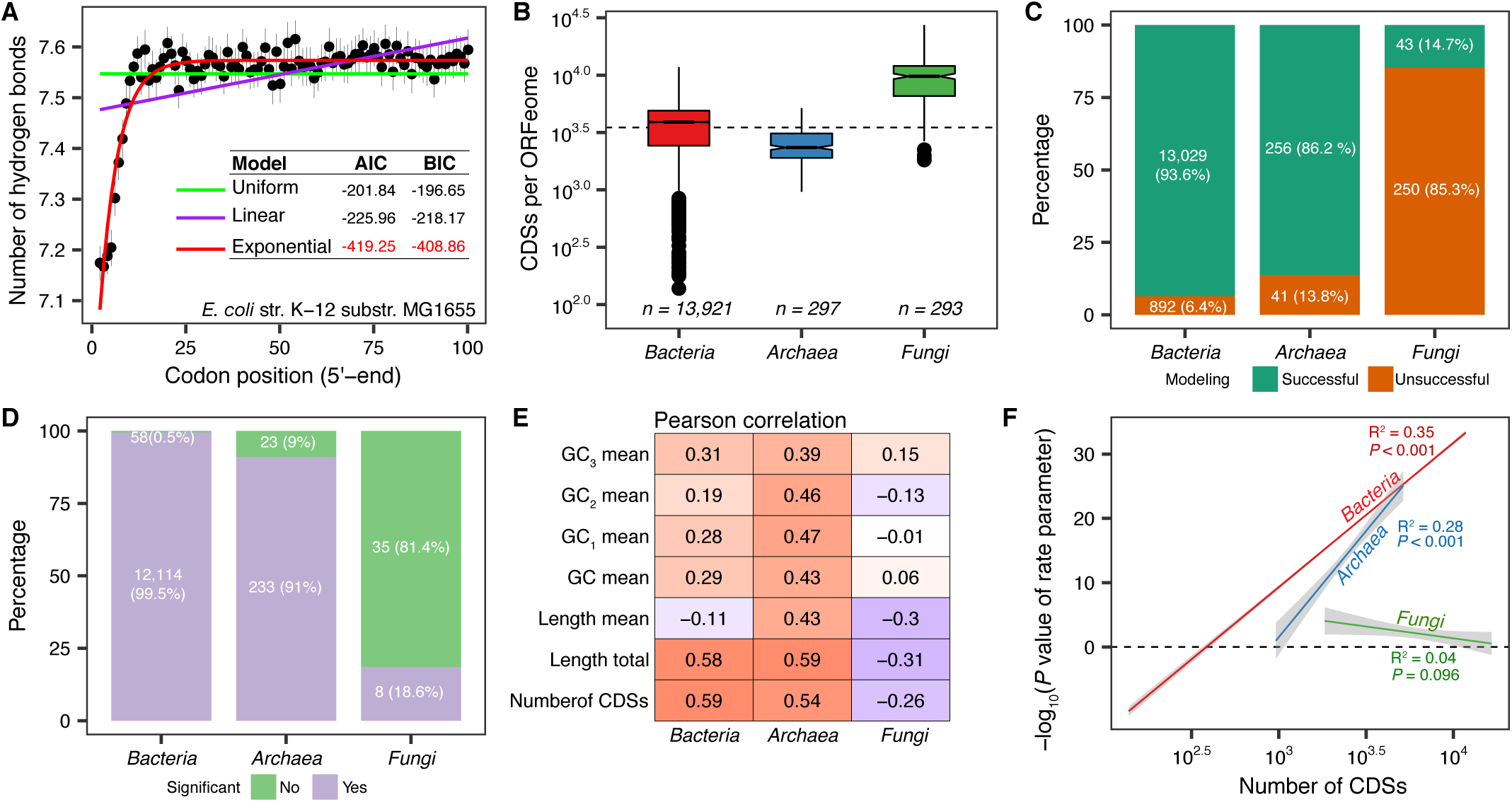
A prokaryotic ramp of the number of hydrogen bonds. **(A)** Three mathematical models were fitted to the hydrogen bonding data. The bounded exponential model with three parameters (red line) produced the best fit to the observed data of *E. coli* ORFeome. Number of CDSs per genome of the datasets used in the analyses (*n* is the number of organisms). **(C)** Percentage of ORFeomes that fitted the bounded exponential model. **(D)** Percentage of ORFeomes that was significant (*P* < 0.001) when modeled with the bounded exponential function. **(E)** Pearson correlation coefficient between the significance of the rate parameter (− *log*_10_ of *P* value) estimated from the bounded exponential model and the different molecular features of ORFeomes analyzed in this study. **(F)** Linear regression model with the number of CDSs per genome as independent variable and the significance of the rate parameter (− *log*_10_ of *P* value) as dependent variable. Grey-shaded region is the 95% confidence interval of the regression model.

We further tested whether the observed ramp in *E. coli* is a conserved feature of ORFeomes in the different domains of life. To investigate this question, we compiled a dataset with ∼14,500 ORFeomes that included *Bacteria* (*n* = 13, 921), *Archaea* (*n* = 297), and *Fungi* (*n* = 293, the representative of eukaryota) (Figure 2B). The dataset comprised ORFeomes with varying total length (Supplemental Figure S2A) and mean CDS length (Supplemental Figure S2B), diverse GC_3_/GC ratio (Supplemental Figure S2C), and organisms from diverse phyla (Supplemental Figure S2D) with multiple mutational biases per phylum (Supplemental Figure S2E). We analyzed the position-dependent number of hydrogen bonds per codon of each ORFeomes and found that in most prokaryotes (94% of *Bacteria* and 86% of *Archaea*), the number of hydrogen bonds per codon position could be successfully fitted by the bounded exponential model whereas the fit was unsuccessful in most *Fungi* (85%) (Figure 2C). Instead, the linear model produced a better fit for most of the fungal ORFeomes (Supplemental Figure S3). We further investigated differences between the successfully and unsuccessfully modeled groups and only two significant different features were observed (Supplemental Figure S4). First, the total ORFeome length tends to be different between the two modeled groups in *Bacteria* and *Fungi* (Supplemental Figure S4A, *P* < 0.001) and second, the mean length of CDS per genome is significantly different in *Bacteria* (Supplemental Figure S4B, *P* < 0.001). No differences were found for GC_3_/GC ratio (Supplemental Figure S4C). When scrutinized by phylum, only *Aquificae* and *Nitrospirae* showed major differences in GC content (Supplemental Figure S4D) and mutational bias (Supplemental Figure S4E) between the two modeled groups (caused by outlier ORFeomes). For the outliers ORFeomes that could not be successfully model, they have a relatively higher GC content and a higher GC_3_/GC ratio. In *Fungi*, the subset of ORFeomes successfully fitted by the bounded exponential model is not monophyletic (Supplemental Figure S5).

Once we established that the bounded exponential model could be fitted to most prokaryotes, we evaluated the statistical significance of the modeling by gathering the *P* value estimated for the rate parameter (a strong indicator of the ramp) in each successful fitted model (Figure 2D). We found that most of the rate parameter estimates for *Bacteria* (99.5%) and *Archaea* (91%) were significant (*P* < 0.001), while only eight were significant in the small subset of ORFeomes that were successfully modeled in *Fungi* (43 ORFeomes) (Figure 2D). We further assessed whether the significance of the rate parameter correlated with other molecular features (Figure 2E). We found that the strongest correlation in prokaryotes was with the total length of the ORFeome and the number of CDSs per ORFeome (Pearson correlation coefficient, Figure 2E). By linear regression modeling, we observed that ∼30% of the variation in the significance of the rate parameter can be explained by the variation in the number of CDSs in the ORFeomes of prokaryotes (*R*^2^ = 0.35 with *P* < 0.001 in *Bacteria* and *R*^2^ = 0.28 with *P* < 0.001 in *Archaea*, Figure 2F).

### Characteristics of the ramp of the number of hydrogen bonds

Significant differences (*α* = 99.9%) were not observed in the estimated parameter of carrying capacity of hydrogen bonds between *Bacteria, Archaea*, and *Fungi* (Figure 3A, adjusted *P* < 0.001). On the other hand, the estimated parameters of initial number of hydrogen bonds (Figure 3B) and rate (Figure 3C) were significantly different between all groups (adjusted *P* < 0.001). We observed that the initial number of hydrogen bonds is the lowest in *Bacteria* (Figure 3B), which is consistent with the rate of increase in the number of hydrogen bonds per codon being the highest in *Bacteria* (Figure 3C) to reach a carrying capacity that is not significantly different between all groups after the ramp (Figure 3A). Hence, by linear regression modeling between the estimated parameters for initial content and carrying capacity, one can approximate how fast is the change in the average number of hydrogen bonds per codon given that the carrying capacity becomes steady at about the 20th codon position (Figure 3D).

**Figure 3.**
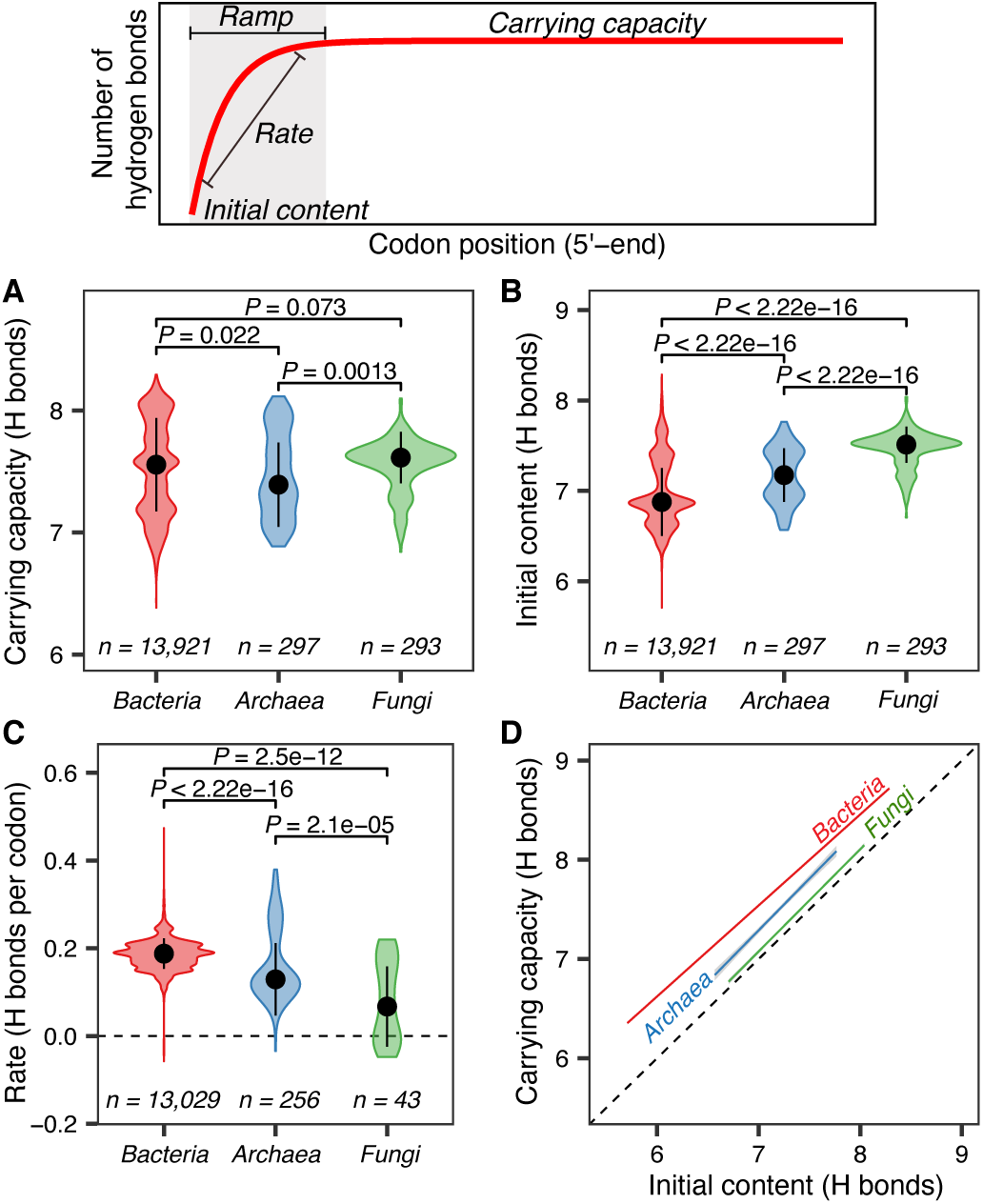
Estimated parameters for the bounded exponential model of the number of hydrogen bonds along CDSs (top panel). In all panels, reported *P* values correspond to the Wilcox test adjusted for multiple testing, and *n* is the number of ORFeomes for which the parameter could be successfully estimated. Comparison of the distribution of estimated **(A)** carrying capacity of hydrogen bonds, **(B)** initial content of the number of hydrogen bonds, and **(C)**rate of the ramp between *Bacteria, Archaea* and *Fungi*. **(D)** Linear regression model with the initial number of hydrogen bonds as independent variable and the carrying capacity of hydrogen bonds as dependent variable.

### Reduced number of hydrogen bonds per codon is selected for at the 5′-end of prokaryotic CDSs

Thus far, we have established that the ramp of hydrogen bonding is conserved in prokaryotes (Figure 2) and that the ramp is characterized by a higher rate of increase in the number of hydrogen bonds per codon in *Bacteria* due to a lower initial number of hydrogen bonds (Figure 3). Based on these findings, we hypothesized that selection acts, through position-dependent codon usage bias, against uniform distribution of hydrogen bonds per codon along CDSs. To test this hypothesis, we compiled a reference set with ∼1,200 bacterial and ~300 archaeal ORFeomes and applied codon shuffling techniques^27,28^ to generate ∼300,000 simulated ORFeomes that contain random synonymous mutations.

The codon-shuffled ORFeomes were used as a null model to test selection against uniformity using the *χ*^2^ statistic^27,28^. The *z*^2^ value (from the *χ*^2^ statistic) per codon position showed that selection acts against uniform distribution of the number of hydrogen bonds and that selection is noticeably stronger at the 5′-end of the *E. coli* ORFeome (Figure 4A). It is also evident that selection against uniformity forms a ramp along the 5′-end of CDSs and that the ramp of selection is conserved in prokaryotic ORFeomes (Figure 4B and Supplemental Figure S6A). Lastly, we investigated the direction of selection acting on the 5′-end of ORFeomes. To assess the selection direction, we computed the value for the *χ*-gram and found that selection acts to reduce the number of hydrogen bonds at the 5′-end of CDSs in the *E. coli* ORFeome following a position-dependent manner (Figure 4C). Similarly, when the analysis was performed on the reference set of ORFeomes, we found that the ramp is conserved in prokaryotes (Figure 4D and Supplemental Figure S6B).

**Figure 4.**
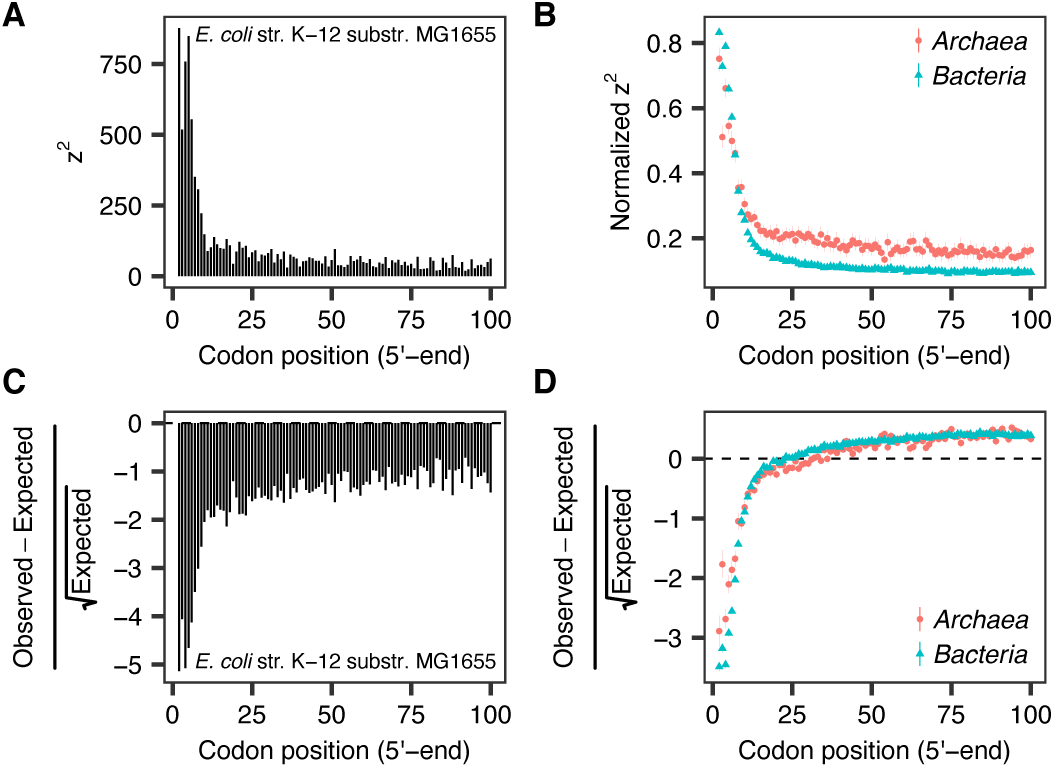
Tests of selection against uniform distribution of the number of hydrogen bonds per codon. **(A)** *z*^2^ value per codon position according to the *χ*^2^ statistic. The higher the *z*^2^ value the higher the selection acting against uniform distribution. **(B)** Mean and the 95% confidence interval of the mean with 1,000 bootstraps of the *z*^2^ value normalized by *min* −*max* normalization (Eq. 4) of all ORFeomes in the representative dataset. **(C)** *χ*-gram value (Eq. 3) per codon position. **(D)** Mean and the 95% confidence interval of the mean with 1,000 bootstraps of the scaled *χ*-gram value (Eq. 3) of all ORFeomes in the representative dataset.

### A web-based application to analyze position-dependent hydrogen bonding

In order to facilitate the analysis of position-dependent hydrogen bonding of novel and custom ORFeomes, a web-based graphical user interface (GUI) application was developed using the `R` package `shiny`^29^. The application incorporates all the methods developed and implemented in this work. In a simple GUI (Supplemental Figure S7), the application allows interactive investigation of novel and customized ORFeomes, download of raw analysis and modeling data, and generation of high-quality figures. The application also reports summary statistics associated with modeling of hydrogen bonding per codon position by the bounded exponential model. For cases that cannot be successfully modeled, the application outputs graphically the observed number of hydrogen bonds per codon position and a summary report of the analysis. The application is publicly available at https://juanvillada.shinyapps.io/hbonds/.

## Discussion

We analyzed over 14,000 bacterial, archaeal, and fungal ORFeomes and found evidence for an exponential ramp of hydrogen bonding at the 5′-end of CDSs in prokaryotes that is created by a position-dependent codon usage bias. With the methods used in this investigation, a similar ramp in fungal ORFeomes was not identified. From a resource allocation perspective, a ramp of hydrogen bonding found in prokaryotes may provide an energy-efficient mechanism to unwind dsDNA and facilitate transcript elongation by reducing the local energy required to melt hydrogen bonds^30–32^. It has been reported previously that AU-rich codons are selected for at the beginning of CDSs in *E. coli*^33^, which would in turn reduce the local hydrogen bonding at the *5*′-end of CDSs. However, a ramp of the number of hydrogen bonds (or a ramp of GC content) is reported for the first time in this study. Different from the approaches applied here, previous studies were limited to characterizing only the first 15 to 20 codon positions^33^. In contrast, we analyzed a longer region of the 5′-end of CDSs (100 or 250 codon positions), which allowed us to identify the formation of the ramp. Besides showing that local hydrogen bonding is reduced and widely conserved at the 5′-end of prokaryotic CDSs, we report a smooth gradual increase of hydrogen bonding that can be modeled by a bounded exponential function.

The most parsimonious explanation for the existence of a ramp of hydrogen bonding in prokaryotes, but not in eukaryotes, is that it is a molecular mechanism that optimizes the coupling of transcription and translation. Transcription and translation in prokaryotes are coupled in space and time^34^ so that variations in one process affect the other. Evolutionary traits may have been developed in order to optimally couple the transcription of protein-coding genes and the translation initiation of mRNA in prokaryotes. A ramp of hydrogen bonding can be one such trait that optimizes transcription efficiency at the 5′-end of CDSs so that transcription can be efficiently coupled with a ramp of translation efficiency found also at the 5′-end of CDSs^11,35,36^.

Although both transcription and translation seem to be mediated by an initial ramp, the ramps have opposite efficiency. While a ramp of translation efficiency has been shown to start with higher occurrence of non-optimally translated codons at the 5′-end as a mechanism to possibly reduce traffic jams of ribosome downstream in translation elongation^5,11,37,38^, a ramp of transcription efficiency found here starts with optimal codons to seemingly reduce the energy required for unwinding dsDNA. Thus, the ramps of transcription and translation efficiency appear complementary in prokaryotes. The complementary ramps would optimize the coupling of transcription and translation by speeding up transcript elongation at the 5′-end of CDSs while simultaneously slowing down translation at the same genetic region. This complementarity of speed can further reduce conflicts between the transcription and translation machineries^39^. From an evolutionary perspective, it will be interesting to further explore whether transcription or translation exerts a stronger selective pressure on local codon usage bias at the 5′-end of OR-Feomes as the data presented here do not allow distinguishing which mechanism drives the selection.

Although we found that the mean rate of increase of the number of hydrogen bonds per codon of prokaryotes is clearly higher than that of eukaryotes, some eukaryotes still showed a non-negligible rate. We hypothesize that this may be signal of a remnant ramp that was lost in eukaryotes with the evolutionary emergence of packaged genomic DNA in the nucleus and further decoupling of transcription and translation. There is evidence in the literature that shows some nuclear sites can still support coupled transcription and translation in eukaryotes^40^.

Overall, we report the existence of a ramp of the number of hydrogen bonds that follows a bounded exponential function at the 5′-end of CDSs in prokaryotes. Optimization of transcription efficiency by reducing hydrogen bonding can be a selective force driving the occurrence of AU-rich codons at the 5′-end of CDSs^33^. The results here suggest that effective coupling of transcription and translation at the 5′-end of CDSs of prokaryotes is achieved by natural evolution via increasing the occurrence of synonymous codons that reduce hydrogen bonding.

## Methods

### Quality control of CDSs

CDSs of genomes were obtained from NCBI/RefSeq^41^. The ORFeome of *Escherichia coli* K-12 substr. MG1655 (acc. number GCF_000005845.2_ASM584v2) was analyzed as a showcase example. For all ORFeomes analyzed in this work, CDSs with lengths not divisible by three and shorter than the number of codons analyzed (100 or 250) were removed from the dataset. The start codon was removed from the dataset before conducting any downstream analyses.

### Quantifying the position-dependent number of hydrogen bonds

DNA sequences were analyzed using the R packages `Biostrings`^42^ and `SeqinR`^43^. Nucleotides in each coding sequence were arranged in a matrix with dimensions equal to the number of CDSs as number of rows and the number of codons analyzed as number of columns. After quality control, all the CDSs in an ORFeome were left-aligned from the 5′-end. The number of hydrogen bonds was computed and stored in a matrix according to the nucleotide base composition of CDSs (Adenine (A) = Thymine (T) = 2; Guanine (G) = Cytosine (C) = 3). The number of hydrogen bonds at each codon position in an ORFeome was computed by calculating the mean and the 95% confidence interval of the mean with nonparametric bootstrapping (1,000 bootstraps) using the `Hmisc` ^44^ package in `R`. Matrix analysis and bootstrapping of thousands of ORFeomes were possible due to parallelization of the computational processes in multiple computer cores using the `R` packages `foreach`^45^, `doParallel` ^46^, and `doSNOW`^47^.

The relative number of hydrogen bonds was calculated as the observed content divided by the maximum number of hydrogen bond per amino acid. The scaled number of hydrogen bonds was calculated by centering and scaling the hydrogen bond contents of codons per amino acid using the scale function in `R`.

### Sequence and genomic analyses

A comprehensive dataset of ORFeomes of prokaryotes (n_*total*_ = 14,218; 13,921 *Bacteria*, 297 *Archaea*) and eukaryotes (293 *Fungi*) were retrieved from NCBI/RefSeq^41^. The commands used to compile the ORFeomes were “*Latest RefSeq*” and “*Exclude anomalous*”. A smaller representative dataset of *Bacteria* ORFeomes was compiled based on a previously curated list that has even representation across phyla^18^. The *Archaea* and *Fungi* ORFeomes were relatively small so all were included in all analyses. The representative prokaryote dataset consisted of 1,496 ORFeomes (1,199 *Bacteria*, 297 *Archaea*). Length, GC content of each CDS, and GC content of each nucleotide position within a codon (GC_1_, GC_2_, and GC_3_) were calculated with `SeqinR`^43^. Taxonomic affiliation of all downloaded ORFeomes was mapped using the `XML` file with the accession numbers of the ORFeomes and the table of lineages of all genomes deposited in NCBI. The table of lineages was generated using `NCBItax2lin` (https://github.com/zyxue/ncbitax2lin) with the NCBI taxonomy database (accessed February 2019). Information regarding the complete and representative ORFeome datasets can be found in Supplemental Table S1 and Supplemental Table S2, respectively.

### Model fitting

The uniform model [*y*(*x*) = *A*], linear model [*y*(*x*) = *Bx* + *C*], and bounded exponential model (Eq. 1) were used to model the mean number of hydrogen bonds per codon as a function of codon position (starting from the 2nd codon position). 

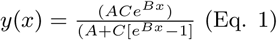

In the models, *y* is the mean number of hydrogen bonds and *x* is the codon position; *A* is the carrying capacity of hydrogen bonds, defined as the maximum average number of hydrogen bonds that a particular codon position can contain in an ORFeome; *B* is the rate of hydrogen bonds per codon, defined as the change in the number of hydrogen bonds per codon; and *C* is the initial content, defined as the number of hydrogen bonds at the first codon after the start codon.

The models were fitted to hydrogen bonding data concerning the first 100 codon positions as the independent variable and the mean number of hydrogen bonds as the dependent variable. Self-Starting Nls Logistic Model was used to estimate the initial parameters and weighted least squares for a nonlinear model was used to estimate the final parameters (both were computed in R). As described previously^27^, the Akaike Information Criterion (AIC) and Bayesian Information Criterion (BIC) were used to select the model that best fitted a dataset. In cases when the exponential model could not be successfully fitted, but parameters were needed for downstream analyses, the initial content and carrying capacity parameters were calculated, respectively, as the minimal number of hydrogen bonds among all codon positions per ORFeome and the trimmed mean number of hydrogen bonds among all codon positions after filtering out 20% of the codons (10 codons from each end).

### Position-dependent null models of ORFeomes with shuffled codons

The null model to test selection against uniform distribution of codons was built by shuffling synonymous codons within all CDSs in each ORFeome. A total of 200 simulated ORFeomes were built for each one of the 1,496 ORFeomes in the representative dataset from which we obtained the metrics of expected and standard deviation of the number of hydrogen bonds per codon position as described in detail elsewhere^28^. Having the observed and expected occurrence of the number of hydrogen bonds per codon, we then computed the *z*^2^ of the *χ*^2^ statistic as shown in (Eq. 2).

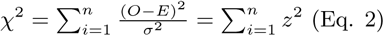

where *O* is the observed count of the number of hydrogen bonds per codon position, *E* is the expected count of the number of hydrogen bonds per codon position computed from the 200 simulated ORFeomes, *σ* is the standard deviation of the number of hydrogen bonds per codon position computed from the 200 simulated ORFeomes, *n* is the number of codon position, and *z* is the *z* score per codon position.

The hanging chi-gram (*χ*-gram) value per position is calculated as shown in Eq. 3. The parameters in Eq. 3 are as defined in Eq. 2. 

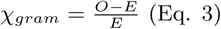

### Statistics, data analysis and data visualization

Data analysis was conducted in `R v3.6.0` using `RStudio v1.2.1335.` The `R` package tidyverse^48^ was used for data analytics, `ggplot2`^49^ for data visualization and `cowplot`^50^ for assembling multiple figure panels. Because sample sizes were large, a *P*-value threshold for significance of 0.001 (*α* = 99.9%) was consistently applied throughout this investigation. Unless otherwise specified, difference between sample groups were tested using two-sided, non-paired Wilcoxon rank sum test (Mann-Whitney test). Correction of *P* values in multiple testing was done with the Benjamini & Yekutieli method. Pearson’s product-moment coefficient was used for linear correlation analyses. Scaled *χ*-gram values were calculated by centering and scaling each ORFeome. Normalized *z*^2^ values were computed using the *min* − *max* normalization function for each ORFeome (Eq. 4). 

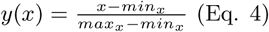

where *x* is the *χ*-gram value (Eq. 3), *min*_*x*_ is the minimum *χ*-gram value of an ORFeome, and *max*_*x*_ is the maximum *χ*-gram value of an ORFeome.

## Data Availability

### Code and data availability

Scripts required to reproduce all the results and figures can be obtained from https://github.com/PLeeLab/H_bonds_ramp. We developed a Web application (https://juanvillada.shinyapps.io/hbonds/) for users to analyze the position-dependent content of hydrogen bonding of ORFeomes.

## Supplementary Data

Supplementary Data are available online.

## Acknowledgements

JCV acknowledges support provided by the Hong Kong PhD Fellowship Scheme (HKPFS).

Author contributions: JCV, MFD and PKHL conceived the study. JCV developed scripts for data analysis. JCV, MFD and PKHL performed data analysis and contributed to the interpretation of findings. JCV and PKHL wrote the manuscript. All authors approved the final version of the manuscript.

## References

[1] J. L. Chaney and P. L. Clark. Roles for synonymous codon usage in protein biogenesis. Annu Rev Biophys, 44:143–66, 2015.

[2] G. Hanson and J. Coller. Codon optimality, bias and usage in translation and mrna decay. Nature Reviews Molecular Cell Biology, 19(1):20–30, 2018.

[3] M. Robinson, R. Lilley, S. Little, J. S. Emtage, G. Yarranton, P. Stephens, M. Millican, M. Eaton, and G. Humphreys. Codon usage can affect efficiency of translation of genes in escherichia coli. Nucleic Acids Research, 12(17):6663–6671, 1984.

[4] E. P. Rocha. Codon usage bias from trna’s point of view: redundancy, specialization, and efficient decoding for translation optimization. Genome Res, 14(11):2279–86, 2004.

[5] E. M. Novoa and L. R. de Pouplana. Speeding with control: codon usage, trnas, and ribosomes. Trends in Genetics, 28(11):574–581, 2012.

[6] J. B. Plotkin and G. Kudla. Synonymous but not the same: the causes and consequences of codon bias. Nature Reviews Genetics, 12(1):32–42, 2011.

[7] Z. E. Sauna and C. Kimchi-Sarfaty. Understanding the contribution of synonymous mutations to human disease. Nature Reviews Genetics, 12(10):683–691, 2011.

[8] G. Cannarrozzi, N. N. Schraudolph, M. Faty, P. von Rohr, M. T. Friberg, A. C. Roth, P. Gonnet, G. Gonnet, and Y. Barral. A role for codon order in translation dynamics. Cell, 141(2):355–367, 2010.

[9] S. Pechmann and J. Frydman. Evolutionary conservation of codon optimality reveals hidden signatures of cotranslational folding. Nature Structural and Molecular Biology, 20(2):237–243, 2013.

[10] Y. M. Zalucki, I. R. Beacham, and M. P. Jennings. Biased codon usage in signal peptides: a role in protein export. Trends in Microbiology, 17(4):146–150, 2009.

[11] T. Tuller, A. Carmi, K. Vestsigian, S. Navon, Y. Dorfan, J. Zaborske, T. Pan, O. Dahan, I. Furman, and Y. Pilpel. An evolutionarily conserved mechanism for controlling the efficiency of protein translation. Cell, 141(2):344–354, 2010.

[12] T. Tuller, Y. Y. Waldman, M. Kupiec, and E. Ruppin. Translation efficiency is determined by both codon bias and folding energy. Proceedings of the National Academy of Sciences of the United States of America, 107(8):3645–3650, 2010.

[13] D. B. Goodman, G. M. Church, and S. Kosuri. Causes and effects of n-terminal codon bias in bacterial genes. Science, 342(6157):475–479, 2013.

[14] S. Pechmann, J. W. Chartron, and J. Frydman. Local slowdown of translation by nonoptimal codons promotes nascent-chain recognition by srp in vivo. Nature Structural and Molecular Biology, 21(12):1100–1105, 2014.

[15] J. O. McInerney. Replicational and transcriptional selection on codon usage in borrelia burgdorferi. Proceedings of the National Academy of Sciences of the United States of America, 95(18):10698–10703, 1998.

[16] X. H. Xia. Maximizing transcription efficiency causes codon usage bias. Genetics, 144(3):1309–1320, 1996.

[17] W. H. Chen, G. T. Lu, P. Bork, S. N. Hu, and M. J. Lercher. Energy efficiency trade-offs drive nucleotide usage in transcribed regions. Nature Communications, 7, 2016.

[18] E. A. Seward and S. Kelly. Selection-driven cost-efficiency optimization of transcripts modulates gene evolutionary rate in bacteria. Genome Biology, 19, 2018.

[19] L. Jeacock, J. Faria, and D. Horn. Codon usage bias controls mrna and protein abundance in trypanosomatids. Elife, 7, 2018.

[20] J. C. Villada, M. F. Duran, and P. K. H. Lee. Genomic evidence for simultaneous optimization of transcription and translation through codon variants in the pmocab operon of type ia methanotrophs. mSystems, 4(4), 2019.

[21] K. S. Murakami and S. A. Darst. Bacterial rna polymerases: the wholo story. Current Opinion in Structural Biology, 13(1):31–39, 2003.

[22] M. Manosas, X. G. Xi, D. Bensimon, and V. Croquette. Active and passive mechanisms of helicases. Nucleic Acids Research, 38(16):5518–5526, 2010.

[23] M. D. Szczelkun and M. S. Dillingham. How to build a dna unwinding machine. Structure, 20(7):1127–1128, 2012.

[24] M. C. Chen, P. Murat, K. Abecassis, A. R. Ferre-D’Amare, and S. Balasubramanian. Insights into the mechanism of a g-quadruplex-unwinding deah-box helicase. Nucleic Acids Research, 43(4):2223–2231, 2015.

[25] A. K. Byrd, D. L. Matlock, D. Bagchi, S. Aarattuthodiyil, D. Harrison, V. Croquette, and K. D. Raney. Dda helicase tightly couples translocation on single-stranded dna to unwinding of duplex dna: Dda is an optimally active helicase. Journal of Molecular Biology, 420(3):141–154, 2012.

[26] M. F. Goodman. Hydrogen bonding revisited: Geometric selection as a principal determinant of dna replication fidelity. Proceedings of the National Academy of Sciences of the United States of America, 94(20):10493–10495, 1997.

[27] A. J. Hockenberry, M. I. Sirer, L. A. N. Amaral, and M. C. Jewett. Quantifying position-dependent codon usage bias. Molecular Biology and Evolution, 31(7):1880–1893, 2014.

[28] J. C. Villada, A. J. B. Brustolini, and W. B. da Silveira. Integrated analysis of individual codon contribution to protein biosynthesis reveals a new approach to improving the basis of rational gene design. DNA Research, 24(4):419–434, 2017.

[29] Chang W, Cheng J, Allaire JJ, Xie Y, and McPherson J. shiny: Web application framework for r. R package version 1. 3. 2, 2019.

[30] W. Ma, K. D. Whitley, Y. R. Chemla, Z. Luthey-Schulten, and K. Schulten. Free-energy simulations reveal molecular mechanism for functional switch of a dna helicase. Elife, 7, 2018.

[31] W. Yang. Lessons learned from uvrd helicase: Mechanism for directional movement. Annual Review of Biophysics, Vol 39, 39:367–385, 2010.

[32] S. S. Patel and I. Donmez. Mechanisms of helicases. Journal of Biological Chemistry, 281(27):18265–18268, 2006.

[33] K. Bentele, P. Saffert, R. Rauscher, Z. Ignatova, and N. Bluthgen. Efficient translation initiation dictates codon usage at gene start. Molecular Systems Biology, 9, 2013.

[34] J. Gowrishankar and R. Harinarayanan. Why is transcription coupled to translation in bacteria? Molecular Microbiology, 54(3):598–603, 2004.

[35] H. Gingold and Y. Pilpel. Determinants of translation efficiency and accuracy. Molecular Systems Biology, 7, 2011.

[36] S. Navon and Y. Pilpel. The role of codon selection in regulation of translation efficiency deduced from synthetic libraries. Genome Biology, 12(2), 2011.

[37] T. Tuller and H. Zur. Multiple roles of the coding sequence 5’ end in gene expression regulation. Nucleic Acids Res, 43(1):13–28, 2015.

[38] J. B. Miller, L. R. Brase, and P. G. Ridge. Extramp: a novel algorithm for extracting the ramp sequence based on the trna adaptation index or relative codon adaptiveness. Nucleic Acids Research, 47(3):1123–1131, 2019.

[39] S. D. Bell and S. P. Jackson. Transcription and translation in archaea: A mosaic of eukaryal and bacterial features. Trends in Microbiology, 6(6):222–228, 1998.

[40] F. J. Iborra, D. A. Jackson, and P. R. Cook. Coupled transcription and translation within nuclei of mammalian cells. Science, 293(5532):1139–1142, 2001.

[41] N. A. O’Leary, M. W. Wright, J. R. Brister, S. Ciufo, D. Haddad, R. McVeigh, B. Rajput, B. Robbertse, B. Smith-White, D. Ako-Adjei, A. Astashyn, A. Badretdin, Y. Bao, O. Blinkova, V. Brover, V. Chetvernin, J. Choi, E. Cox, O. Ermolaeva, C. M. Farrell, T. Goldfarb, T. Gupta, D. Haft, E. Hatcher, W. Hlavina, V. S. Joardar, V. K. Kodali, W. Li, D. Maglott, P. Masterson, K. M. McGarvey, M. R. Murphy, K. O’Neill, S. Pujar, S. H. Rangwala, D. Rausch, L. D. Riddick, C. Schoch, A. Shkeda, S. S. Storz, H. Sun, F. Thibaud-Nissen, I. Tolstoy, R. E. Tully, A. R. Vatsan, C. Wallin, D. Webb, W. Wu, M. J. Landrum, A. Kimchi, T. Tatusova, M. DiCuccio, P. Kitts, T. D. Murphy, and K. D. Pruitt. Reference sequence (refseq) database at ncbi: current status, taxonomic expansion, and functional annotation. Nucleic Acids Res, 44(D1):D733–45, 2016.

[42] H. Pagès, P. Aboyoun, Gentleman R., and DebRoy S. Biostrings: Efficient manipulation of biological strings. R package version 2.52.0, 2019.

[43] D. Charif, J. Thioulouse, J. R. Lobry, and G. Perriere. Online synonymous codon usage analyses with the ade4 and seqinr packages. Bioinformatics, 21(4):545–547, 2005.

[44] F. E. Harrell. Hmisc: Harrell miscellaneous. R package version 4.2-0, 2019.

[45] S. Weston, H. Ooi, and Microsoft. foreach: Provides foreach looping construct for r. R package version 1.4.4, 2017.

[46] S. Weston and Microsoft. doparallel: Foreach parallel adaptor for the ‘parallel’ package. R package version 1.0.14, 2018.

[47] S. Weston and Microsoft. dosnow: Foreach parallel adaptor for the ‘snow’ package. R package version 1.0.16, 2017.

[48] H. Wickham. tidyverse: Easily install and load the ‘tidyverse’. R package version 1.2.1, 2017.

[49] H. Wickham. ggplot2: Elegant Graphics for Data Analysis. Use R! Springer, New York, 2016.

[50] C. O. Wilke. cowplot: Streamlined plot theme and plot annotations for ‘ggplot2’. R package version 1.0.0, 2019.

