## Supplementary Data for "Codon usage bias creates a ramp of hydrogen bonding at the 5′-end in prokaryotic ORFeomes"

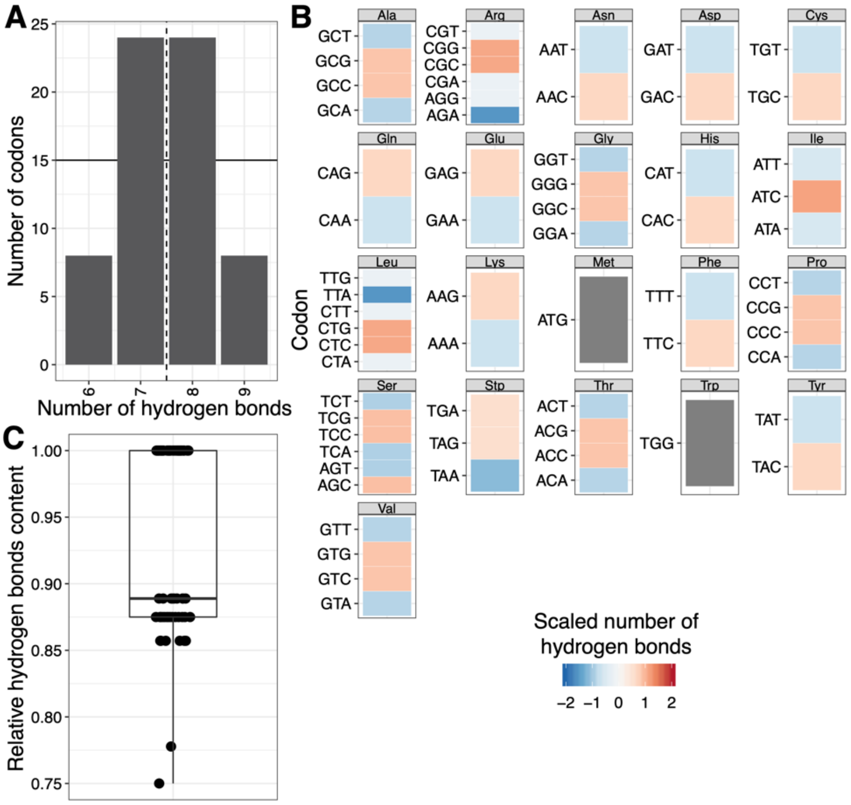


**Figure S1.** Optimization space for the number of hydrogen bonds as a function of codon usage. **(A)** Frequency of codons according to the number of hydrogen bonds a codon can contain. **(B)** The number of hydrogen bonds (scaled value is shown) for each amino acid by codon. Synonymous codon choices can reduce or increase the number of hydrogen bonds of each amino acid. Scaled values were calculated by centering and scaling the number of hydrogen bonds of codons that code for the same amino acid. **(C)** The hydrogen bonding content of codons relative to the maximum possible content among synonymous codons. Relative content was calculated as the number of hydrogen bonds of each codon divided the maximum number of hydrogen bonds per amino acid.

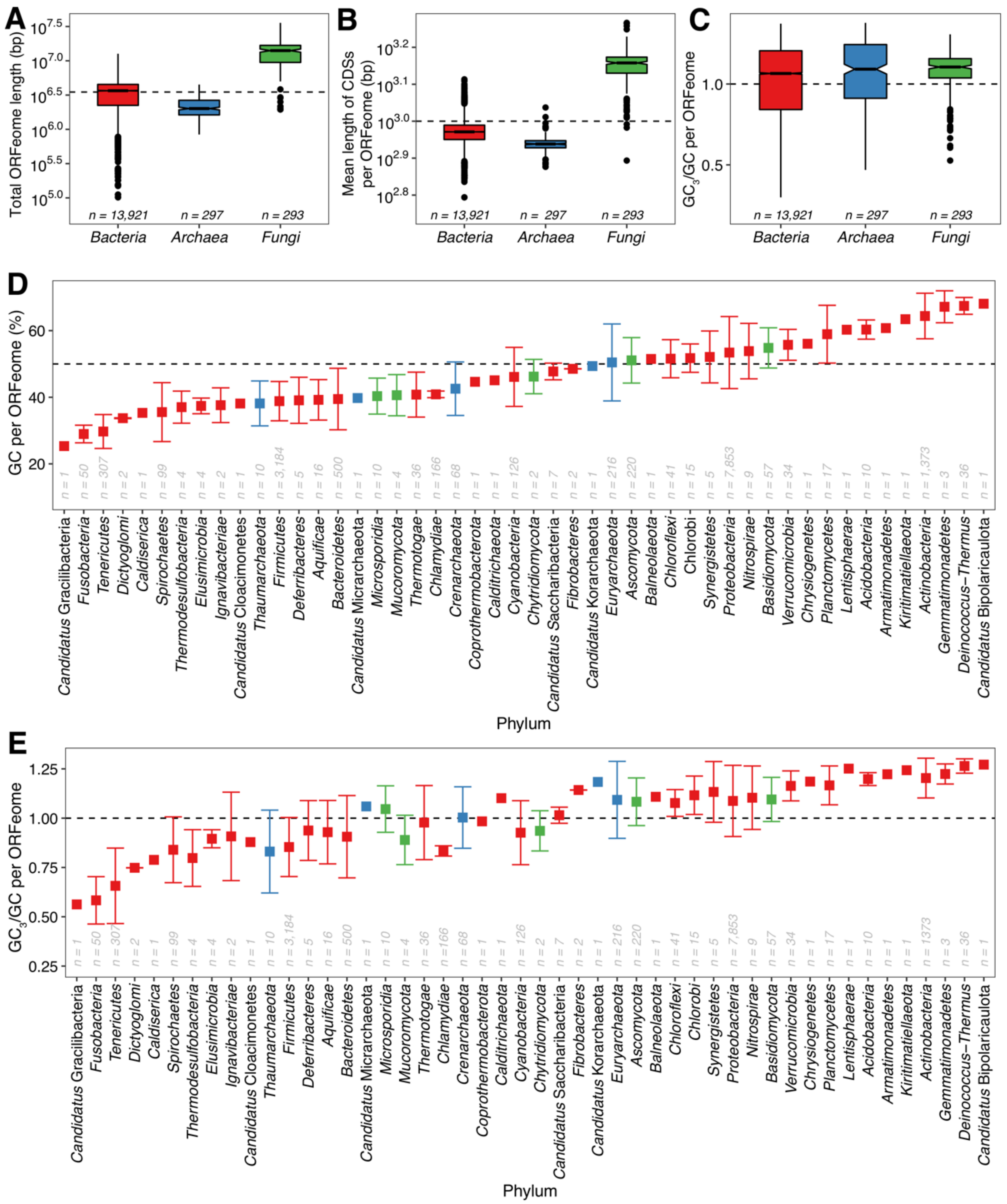


**Figure S2.** Characteristics of ORFeomes analyzed in this study. **(A)** Total ORFeome length (in base pair, bp), **(B)** mean length of CDSs in each genome, and **(C)** distribution of mutational biases in each ORFeome computed as the ratio of $GC_3$ to GC content in *Bacteria*, *Archaea* and *Fungi*. **(D)** Mean GC content and **(E)** mean $GC_3$ to GC content ratio of all CDSs in each ORFeome grouped by phyla. Error bars represent one standard deviation.

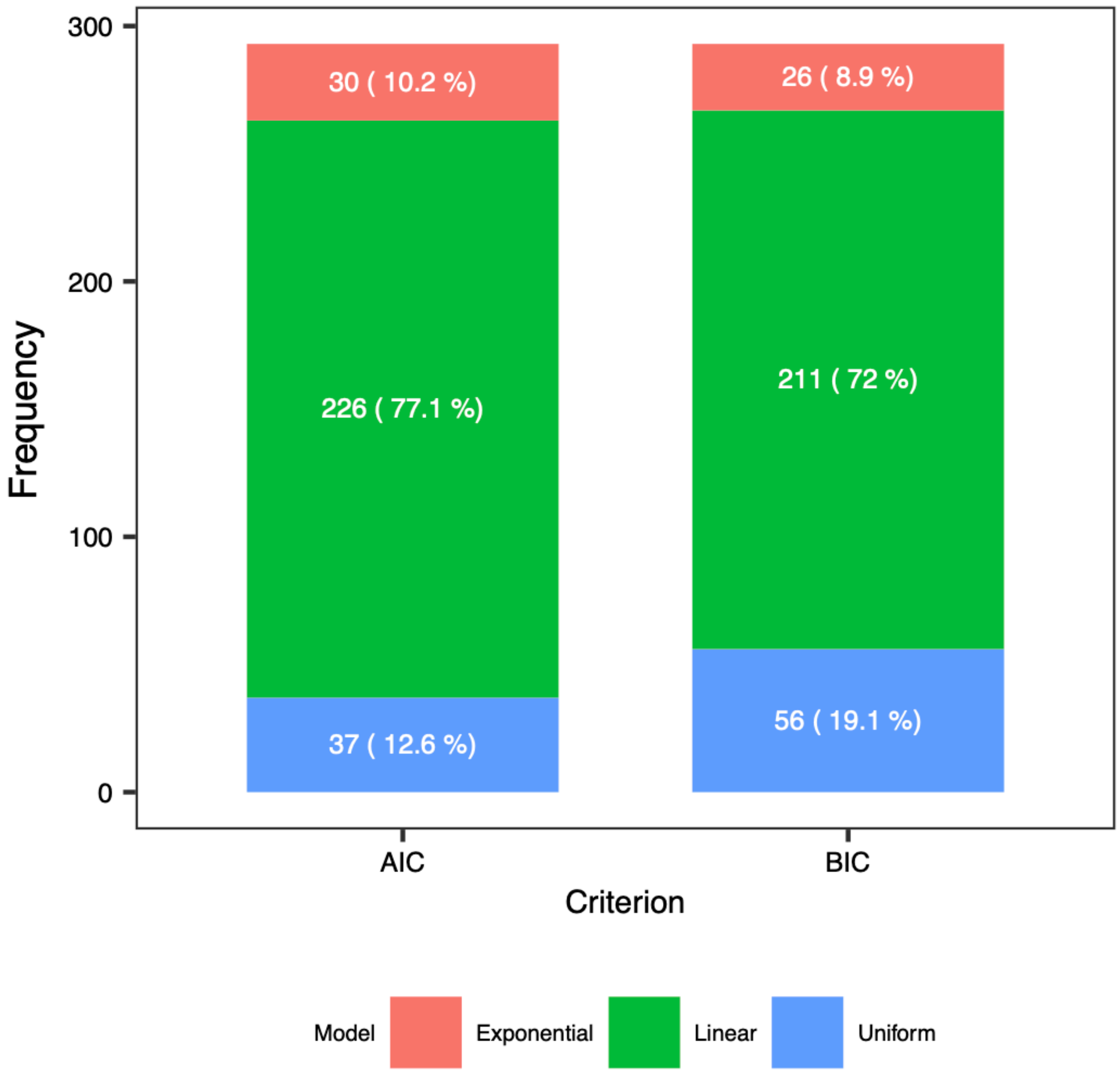


**Figure S3.** Frequency of the best fitted model in fungal ORFeomes (*n* = 293) according to the Akaike Information Criterion (AIC) and Bayesian Information Criterion (BIC).

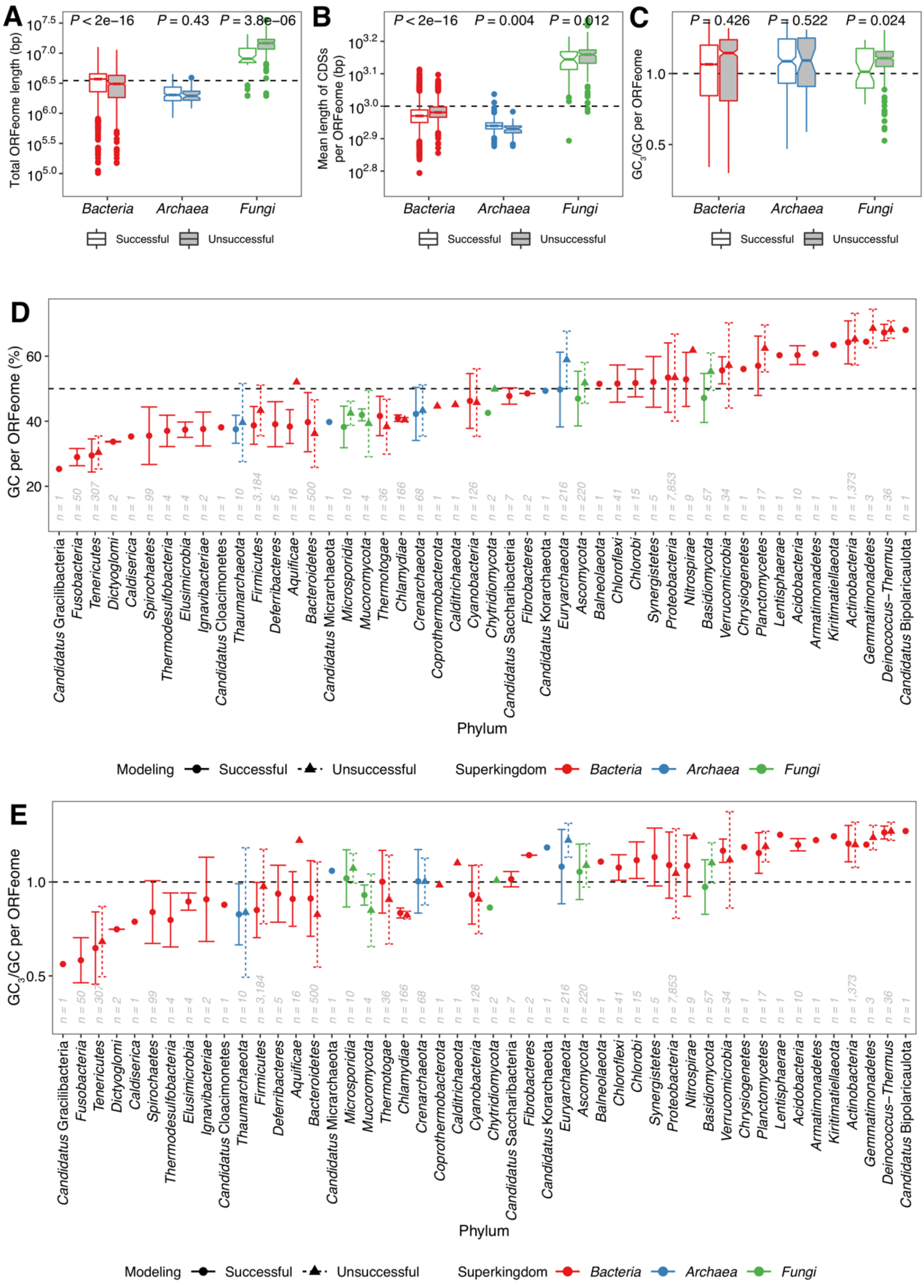


**Figure S4.** Characteristics of ORFeomes analyzed in this study divided into two groups based on whether modeling of the number of hydrogen bonds per codon position was successfully or unsuccessfully fitted by the bounded exponential model. **(A)** Total ORFeome length (in base pair, bp), **(B)** mean length (in bp) of CDSs in each ORFeome, and **(C)** distribution of mutational biases found in each ORFeome computed as the ratio of $GC_3$ to GC content in *Bacteria*, *Archaea* and *Fungi*. All the indicated *P* values correspond to hypothesis testing of the difference between the successful and unsuccessful groups with the two-tailed unpaired *t*-test. **(D)** Mean GC content and **(E)** mean $GC_3$ to GC content ratio of all CDSs in each ORFeome grouped by phyla. Error bars represent one standard deviation.

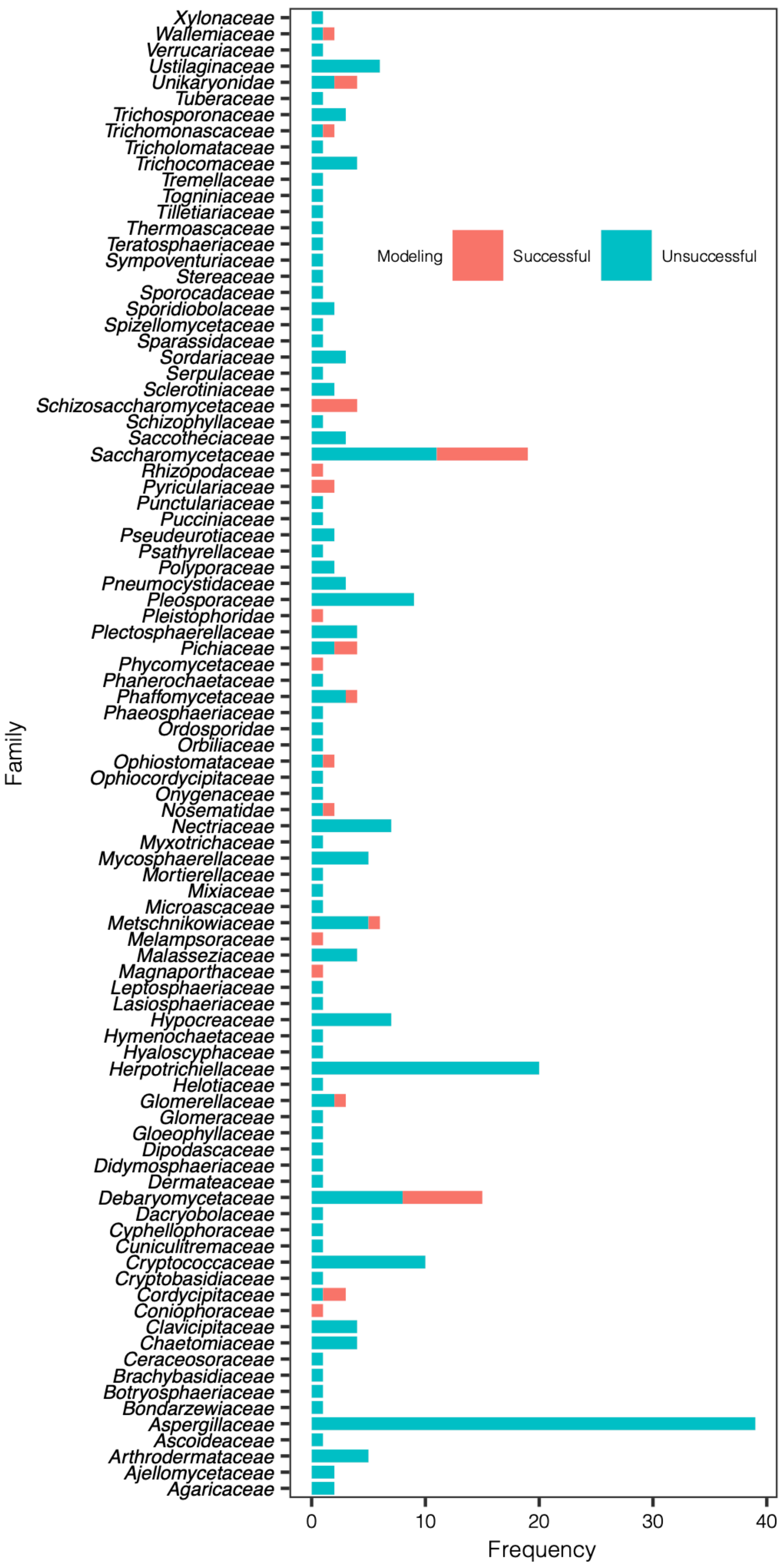


**Figure S5.** Families of fungal ORFeomes successfully and unsuccessfully fitted by the bounded exponential model.

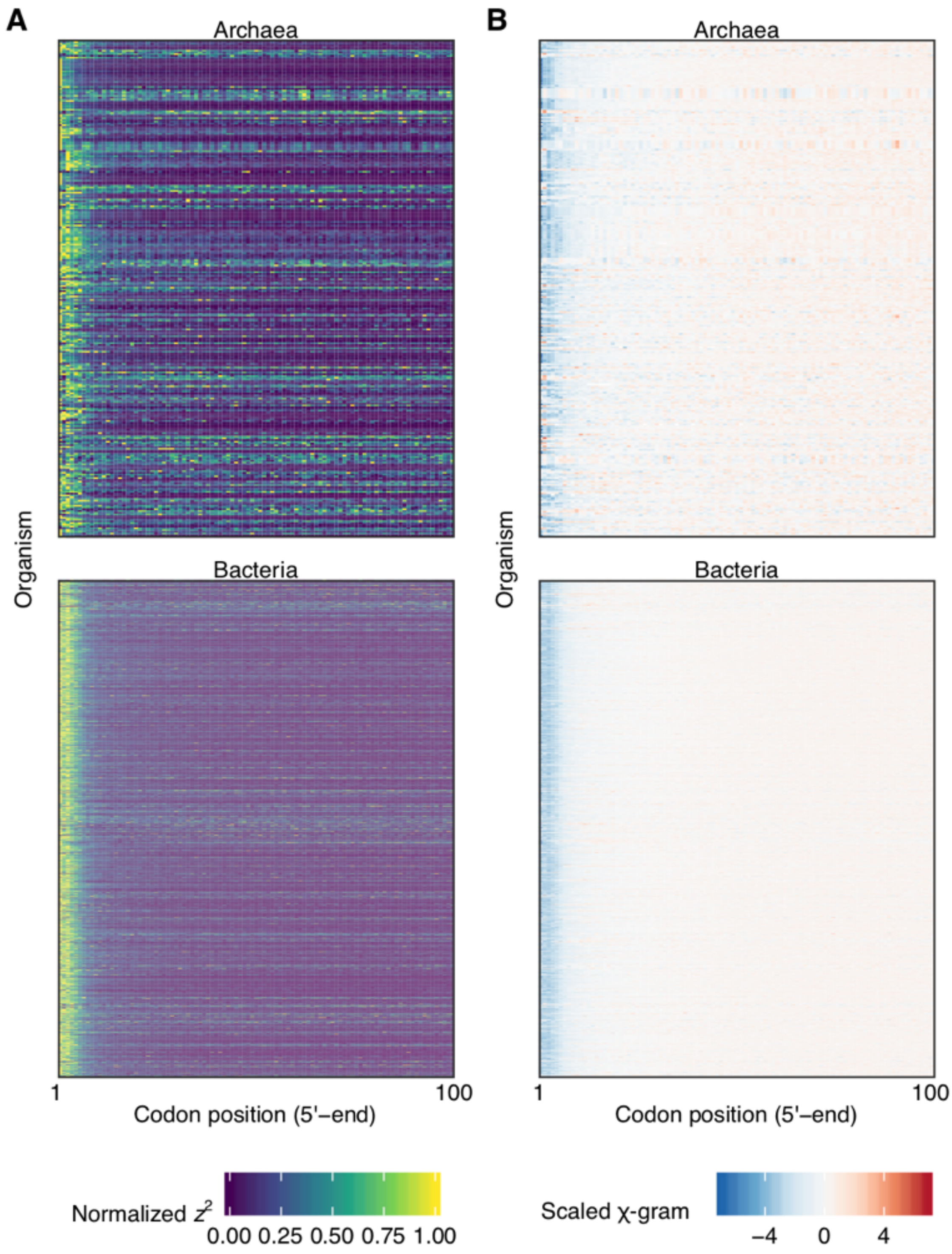


**Figure S6.** Detailed illustration of the ramp of selection against uniform distribution of the number of hydrogen bonds per codon. **(A)** Normalized $z^2$ and **(B)** scaled χ-gram of each ORFeome (~1,500 in total). Columns correspond to the codon position (from

$2^{nd}$ to $100^{th}$) and rows correspond to individual organisms in the representative dataset. The 5′-end (first ~15 codon positions) of CDSs shows the highest selection.

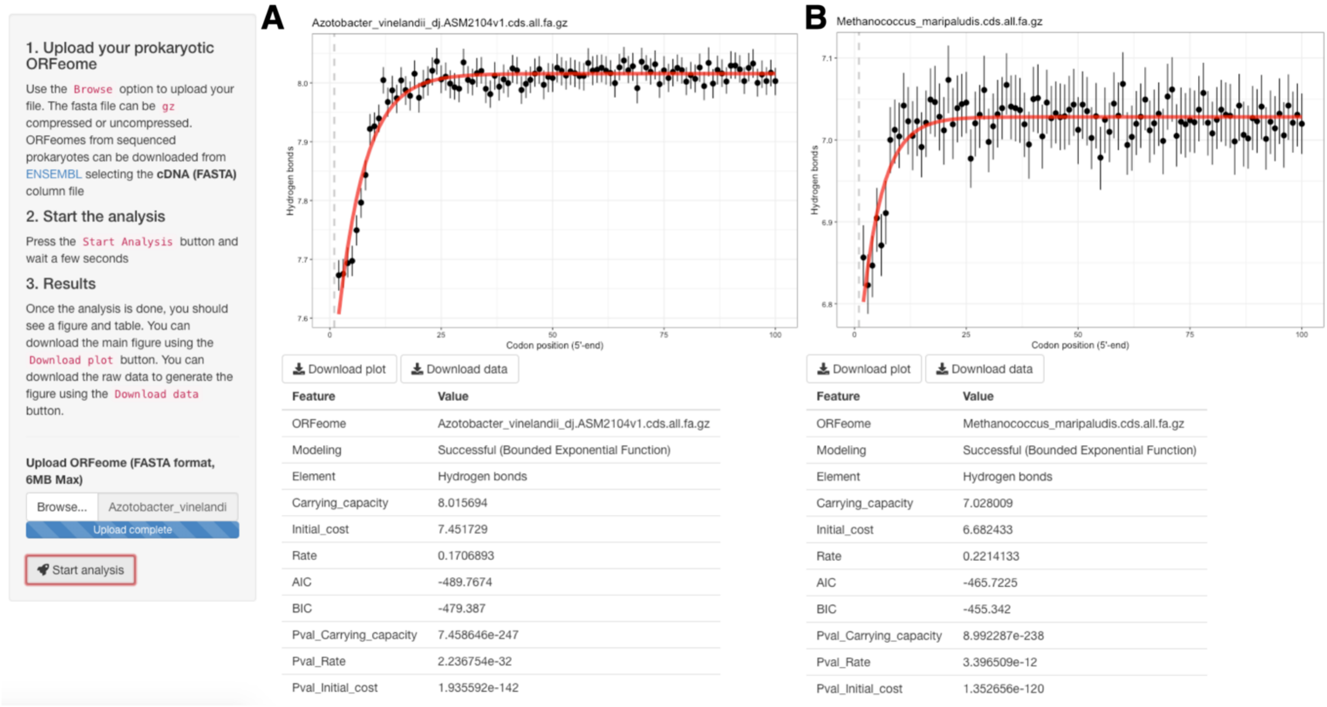


**Figure S7.** Screenshot illustration of the web-based graphical user interface application developed to analyze novel and customized ORFeomes. Examples of analyzing ORFeomes belonging to **(A)** *Bacteria* and **(B)** *Archaea*.

**Table S1.** Information of the comprehensive dataset of ORFeomes analyzed in this study.

**Table S2.** Information of the representative dataset of ORFeomes analyzed in this study.
